# Identification of powdery mildew resistance QTL in *Fragaria x ananassa*

**DOI:** 10.1101/196485

**Authors:** Helen M. Cockerton, Robert J. Vickerstaff, Amanda Karlström, Fiona Wilson, Maria Sobczyk, Joe Q. He, Daniel J. Sargent, Andy J. Passey, Kirsty J. McLeary, Katalin Pakozdi, Nicola Harrison, Maria Lumbreras-Martinez, Laima Antanaviciute, David W. Simpson, Richard J. Harrison

**Author notes:** **Corresponding author:** Richard J. Harrison, **Email:**.

## Abstract

The obligate biotrophic fungus *Podosphaera aphanis* is the causative agent of powdery mildew on cultivated strawberry (*Fragaria x ananassa*). Genotypes from two bi-parental mapping populations ‘Emily’ x ‘Fenella’ and ‘Redgauntlet’ x ‘Hapil’ were phenotyped for powdery mildew disease severity in a series of field trials. Here we report multiple QTL associated with resistance to powdery mildew, identified in ten phenotyping events conducted across different years and locations. Seven QTL show a level of stable resistance across multiple phenotyping events however many other QTL were represented in a single phenotyping event and therefore must be considered transient. One of the identified QTL was closely linked to an associated resistance gene across the wider germplasm. Furthermore, a preliminary association analysis identified a novel conserved locus for further investigation. Our data suggests that resistance is highly complex and that multiple additive sources of quantitative resistance to powdery mildew exist across strawberry germplasm. Implementation of the reported markers in marker-assisted breeding or genomic selection would lead to improved powdery mildew resistant strawberry cultivars, particularly where the studied parents, progeny and close pedigree material are included in breeding germplasm.

**Key Message:** Powdery mildew resistance in two strawberry mapping populations is controlled by both stable and transient novel QTL of moderate effect. Some transferability of QTL across wider germplasm was observed.

## Introduction

*Podosphaera aphanis* (syn. *Sphaerotheca macularis*) is a global pathogen on strawberry (*Fragaria x ananassa*) (Peries 1962), where late season infestations in untreated fields result in unmarketable fruit and severe yield loss (Nelson et al 1995). Powdery mildew was rated the most important disease by large UK strawberry producers (Calleja 2011), with 65% of UK growers reporting common outbreaks of *P. aphanis* (Calleja 2011). The transfer of strawberry field production into protected systems has been associated with a heightened incidence of powdery mildew and greater fungal biomass has been observed on strawberry fruits in polytunnel environments (Xiao et al 2001).

*Podosphaera aphanis* infects the leaves, fruit, stolon and flowers of strawberry plants (Paulus 1990). A higher disease level on the abaxial (lower) leaf surface, is due to infection of emergent susceptible leaves before unfolding, with ontogenic resistance developing in the adaxial surface prior to exposure (Asalf et al 2014). For conidia on host tissue, optimum conditions for germination and colony establishment can lead to disease symptom development within four days, upon which conidiation begins anew (Amsalem et al 2006). Although a high relative humidity (RH) is required for germination and release of conidia (Amsalem et al 2006), conidial germination is inhibited by free water (Peries 1962). The sexual ascospores overwintering on dead plant foliage are considered to be a major sources of infection in early spring, as such, the removal of old strawberry foliage as a source of inoculum should reduce epidemics (Xu et al 2008a).

*P. aphanis* is considered to have a small host range with host specificity of strawberry and raspberry (Harvey and Xu 2010). Across the powdery mildews it is understood that many host-specific adaptations have arisen through convergent evolution, with over 400 different fungal species causing powdery mildew on 9838 different angiosperm hosts (Amano 1986; Braun 1987; Mori et al 2000).

For strawberries, powdery mildew is primarily controlled using fungicide application as many varieties have poor levels of disease resistance. Fungicides with modes of action targeting fungal respiration, nucleic acid synthesis, sterol biosynthesis and signal transduction are commonly used to control *P. aphanis* on strawberry (Lainsbury 2016). However, the evolution of resistance to sterol demethylation inhibitor fungicides has posed challenges for *P. aphanis* control (Sombardier et al 2010). Such challenges have been exacerbated by the loss of active ingredients associated with stricter European regulations (e.g. 91/414/EEC; (Colla et al 2012), highlighting a greater requirement for *P. aphanis* resistant breeding resources.

Previous studies have shown high variation in powdery mildew resistance within strawberry breeding germplasm and high heritability of resistance, indicating the large potential for enhancing disease resistance through breeding (Nelson et al 1995). Utilisation of pre-breeding data and marker-assisted or genomic selection (Whitaker et al 2012) will aid the production of durable powdery mildew resistance and reduce growers reliance on fungicide control.

## Materials and Methods

Plant material was created through a cross between the powdery mildew resistant strawberry cultivar ‘Emily’ and the susceptible cultivar ‘Fenella’ to produce a mapping population of 181 individuals which segregates for mildew resistance. This was phenotyped over four years in six locations denoted; 2011, 2012a, 2012b, 2013a, 2013b, 2014. All phenotyping events were conducted at East Malling Research, Kent, UK (now NIAB EMR) except 2013b which was conducted in Paraje Moriteja, Rociana del Condado, Spain. An additional pre-established mapping population was phenotyped for mildew resistance; the ‘Redgauntlet’ x ‘Hapil’ (RxH) mapping population (168 individuals) was phenotyped in 2012, 2013, 2014 and 2016 at East Malling (Sargent et al 2012). Plants were maintained in a polytunnel and runners were pinned down into 9 cm pots containing compost and transferred into polythene covered raised beds with trickle irrigation. Raised beds were fumigated with chloropicrin to control soil borne pests and diseases. Plants were strimmed in early July (or mid April in Spain; 2013b) to remove old leaf material and expose young foliage; strimming ensures simultaneous disease development of new leaf material. Plantings were downwind of established strawberry plots, which provided a natural source of inoculum. Infection of mildew was allowed to establish within the field plots. Plants were arranged in a randomized block design with 3-6 replicate plants per genotype. Disease scores were recorded twice between late July and early September in UK field plots as dictated by disease symptom progression, and during May in the Spanish plot. Plants were scored for mildew disease symptoms based on an existing scale (Simpson 1987), where scores denote: 1- a healthy plant with no visible disease symptoms, 2- slight leaf curling with no visible mycelium, 3- leaf curling and mottling 4-severe leaf curling, redding and visible damage to lower leaf surface, 5- severe necrosis and some leaf death. A validation set of 75 cultivars and accessions were phenotyped in 2017 with 10 replicate plants per accession.

### Linkage map generation

DNA was extracted from new leaf material using the Qiagen DNAeasy plant mini extraction kit according to the manufacturer’s instructions. Biparental populations were genotyped using the Affymetrix Istraw90 Axiom^®^ array (i90k) containing approximately 90 thousand potential genetic markers (Bassil et al 2015). Cultivars and accessions were genotyped on either the i90k and/or the streamlined Axiom^®^ IStraw35 384HT array (i35K, containing approximately 35 thousand markers; Verma et al 2017). The linkage maps were created using the Crosslink program (https://github.com/eastmallingresearch/crosslink) designed for octoploid linkage map development (Vickerstaff and Harrison 2017). Haplotype blocks lacking recombination in any of the progeny were identified for each mapping population and were used to identify neighbouring SNPs identified in both mapping populations that may represent the same QTL and resistance allele. In the ‘Redgauntlet’ x ‘Hapil’ linkage map the average distance between markers is 0.75 cM, there are gaps > 20 cM on chromosome 1D, 4D and 6C. In the ‘Emily’ x ‘Fenella” linkage map the average distance between markers is 0.71 cM there are gaps >20 cM on chromosome 2C, 3B, 3D, 4D and 5C.

## Statistical analysis

### Phenotype calculation

The area under the disease progression curve (AUDPC) was calculated for each phenotyping event using the R package “agricolae” (Felipe 2017) to predict scores for QTL analysis. Best linear unbiased prediction (BLUP) was calculated using the relative AUDPC across phenotyping events, to predict overall genotype scores for QTL analysis using the R package “nlme” (Pinheiro et al 2017).

### QTL identification

Disease resistance QTL were identified using Kruskal–Wallis analysis for each marker; the most significant marker was automatically selected for each linkage group and marker type before conducting a stepwise AIC linear model selection in R. QTL effect size was calculated based on the output parameters from the predictive linear model. A permutation test was conducted based on 10,000 iterations of randomly selected data to determine that significance selection value was not required. The consensus map was generated using marker data from five mapping populations (‘Redgauntlet’ x ‘Hapil’, ‘Emily’ x ‘Fenella’, ‘Flamenco’ x ‘Chandler’, ‘Capitola’ x ‘CF1116’; INRA, ‘Camerosa’ x ‘Dover’; CRAG). Marker positions were anchored to the *F. vesca* genome v2.0 (Tennessen et al. 2014), to allow the locations of QTL to be placed onto the physical map.

### Comparison of phenotyping events

Mixed effect models using the relative AUDPC values were used to assess the relative importance of genotype, environment and Genotype x Environment interactions on disease severity across phenotyping events. The models with and without each component were compared using analysis of variation (ANOVA). Normal residuals for AUDPC scores were confirmed using the Kolmogorov-Smirnov test. A two-way ANOVA on the relative AUDPC values allowed comparison of disease severity between phenotyping events.

### QTL validation

The QTL analysis is conducted using the i90k marker data in order to best represent the position of the resistance marker, however the QTL analysis was repeated using the subset of i90k markers represented in the i35k chip and validation set, allowing the identification of substitute i35k markers associated with each QTL and comparison with an expanded validation panel of cultivars. Substitute i35k markers are on the same haplotype as the QTL identified in the i90k analysis in the mapping population, therefore they are analogous but may have a weaker association with resistance due to their increase genetic distance from the ‘best’ marker in the mapping population. Restricted maximum likelihood (REML) was used to determine the strength of association between the substitute focal SNPs and the resistance allele in the wider germplasm using the R package “lme4” (Bates et al 2014) allowing the identification of markers in linkage disequilibrium (LD) with the QTL. A linear model between the observed and predicted phenotype was produced for QTL with strong marker-trait associations.

### Association study

The subset of SNPs present on the Istraw90k chip, showing at least 10 % minor allele frequency were screened for association with mildew resistance (https://github.com/harrisonlab/popgen/blob/master/snp/gwas_quantitative_pipeline.md). This association analysis was conducted on 75 validation accessions using Plink (Purcell et al 2007), *p* values were corrected for population structure and adjusted using the Benjamini-Hochberg multiple test correction

### Identification of candidate resistance genes

NB-LRR, TM–CC, RLP, RLK (S-type and general) were identified within the *F. vesca* genome (assembly v1.1) (Shulaev et al 2011) by screening gene models for motifs following established pipelines (Li et al 2016). Candidate susceptibility factors candidate MLO genes identified in Rosaceous crops by Pessina et al. (Pessina et al 2014) and resistance genes were identified within 100 kbp of the significant QTL using BEDtools (Quinlan and Hall 2010) and tblastx (Karlin and Altschul 1993) against the NCBI database to determine any characterised function of homologous genes.

## Results

### Disease pressure varies between years and sites

The greatest proportion of variance in the relative AUDPC values was explained by the environment for both populations (‘Emily’ x ‘Fenella’: *X*^*2*^_(4,5;1)_= 1264.7; *p* < 0.001, ‘Redgauntlet’ x ‘Hapil’: *X*^*2*^_(4,5;1)_ = 374.83; *p* < 0.001) followed by genotype (‘Emily’ x ‘Fenella’: *X*^*2*^_(1)_ = 322.97; *p* < 0.001, ‘Redgauntlet’ x ‘Hapil’: *X*^*2*^_(4,5;1)_ = 143.96; *p* < 0.001). The phenotyping events in 2014 showed low disease symptoms across both populations suggesting either low disease pressure or low environmental conductivity (Fig 1). A significant effect of Genotype x Environment interaction was observed in both populations (‘Emily’ x ‘Fenella’: *X*^*2*^_(4,5;1)_ = 170.66; *p* < 0.001, ‘Redgauntlet’ x ‘Hapil’: *X*^*2*^_(4,5;1)_ = 29.11; *p* < 0.001). Across phenotyping events, broad sense heritability factors varied between 24.1-59.0 for ‘Emily’ x ‘Fenella’ and 40.1- 53.8 for ‘Redgauntlet’ x ‘Hapil’, revealing a moderate proportion of the variation in the data can be explained by the genetic variation, and that there is a moderate to large environmental influence on disease symptom expression (Table 1). Nonetheless, the correlation analysis showed significant, positive correlations between disease scores for all the phenotyping events (Fig 2). The 2013b phenotyping event showed the greatest correlation across all ‘Emily’ x ‘Fenella’ phenotyping events with an average correlation coefficient of 0.58.

**Fig. 1.**
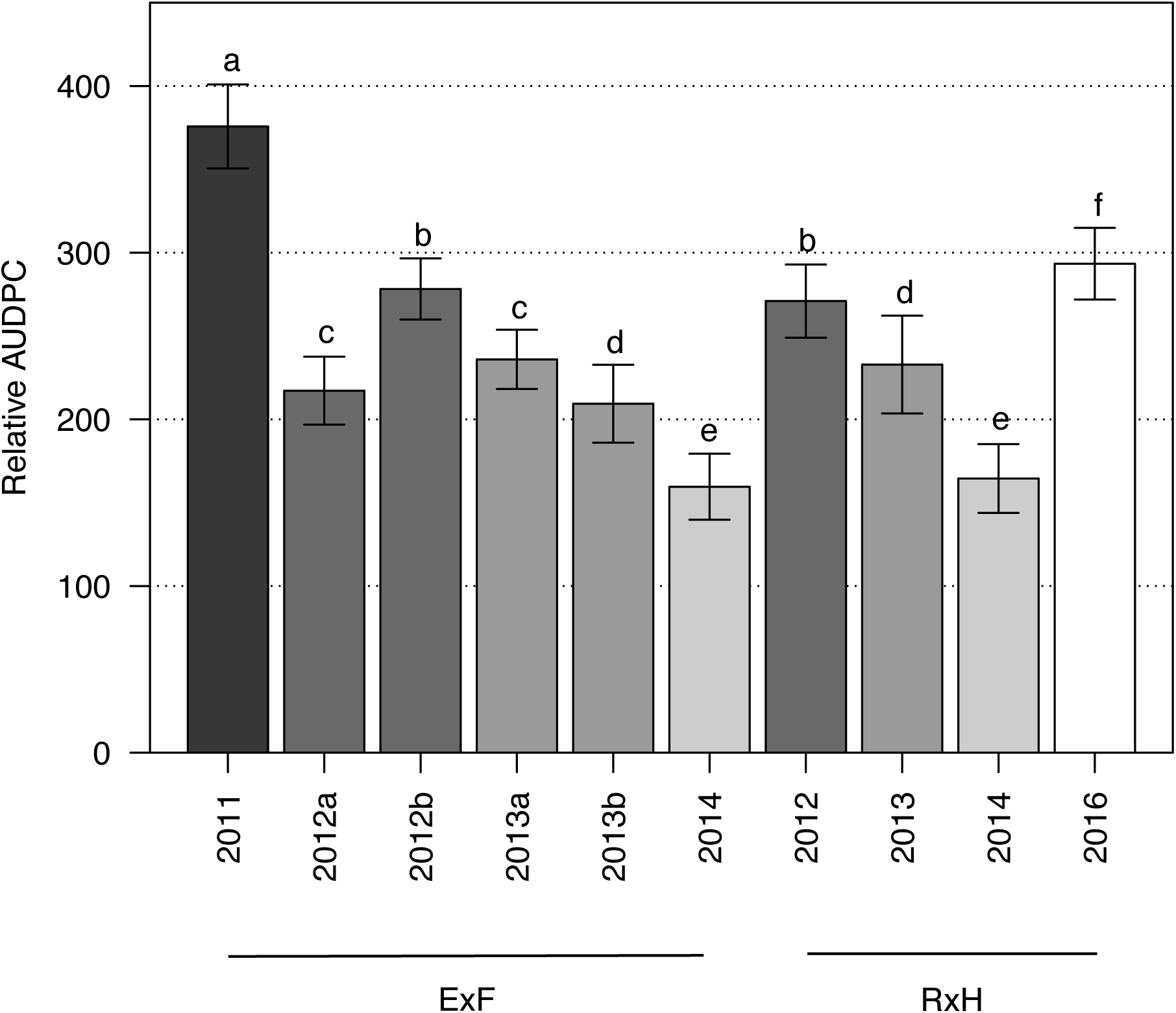
Average relative area under the disease progression curve (rAUDPC) across all genotypes for each phenotyping event. Error bars are standard errors.

**Table 1.**
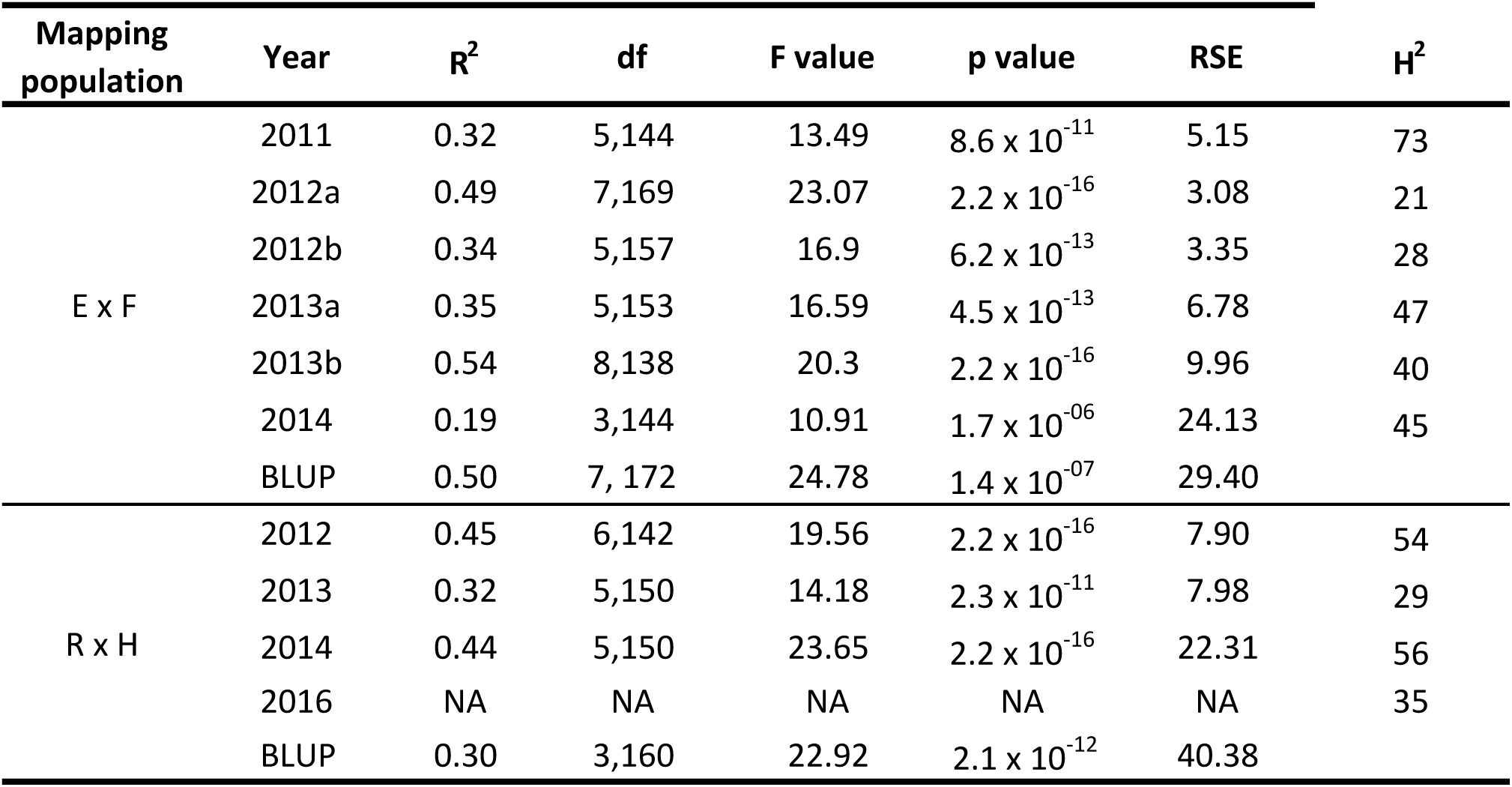
Model parameters for the predictive linear model for each phenotyping event. Predicted versus observed disease scores for each genotype within the population. R^2^ is the coefficient of determination; H^2^ is broad-sense heritability associated with each phenotyping event.

| Mapping population | Year | $R^2$ | df | F value | p value | RSE | $H^2$ |
| --- | --- | --- | --- | --- | --- | --- | --- |
| E x F | 2011 | 0.32 | 5,144 | 13.49 | $8.6 \times 10^{-11}$ | 5.15 | 73 |
| | 2012a | 0.49 | 7,169 | 23.07 | $2.2 \times 10^{-16}$ | 3.08 | 21 |
| | 2012b | 0.34 | 5,157 | 16.9 | $6.2 \times 10^{-13}$ | 3.35 | 28 |
| | 2013a | 0.35 | 5,153 | 16.59 | $4.5 \times 10^{-13}$ | 6.78 | 47 |
| | 2013b | 0.54 | 8,138 | 20.3 | $2.2 \times 10^{-16}$ | 9.96 | 40 |
| | 2014 | 0.19 | 3,144 | 10.91 | $1.7 \times 10^{-06}$ | 24.13 | 45 |
| | BLUP | 0.50 | 7, 172 | 24.78 | $1.4 \times 10^{-07}$ | 29.40 | |
| R x H | 2012 | 0.45 | 6,142 | 19.56 | $2.2 \times 10^{-16}$ | 7.90 | 54 |
| | 2013 | 0.32 | 5,150 | 14.18 | $2.3 \times 10^{-11}$ | 7.98 | 29 |
| | 2014 | 0.44 | 5,150 | 23.65 | $2.2 \times 10^{-16}$ | 22.31 | 56 |
|  | 2016 | NA | NA | NA | NA | NA | 35 |
| | BLUP | 0.30 | 3,160 | 22.92 | $2.1 \times 10^{-12}$ | 40.38 | |

**Fig. 2.**
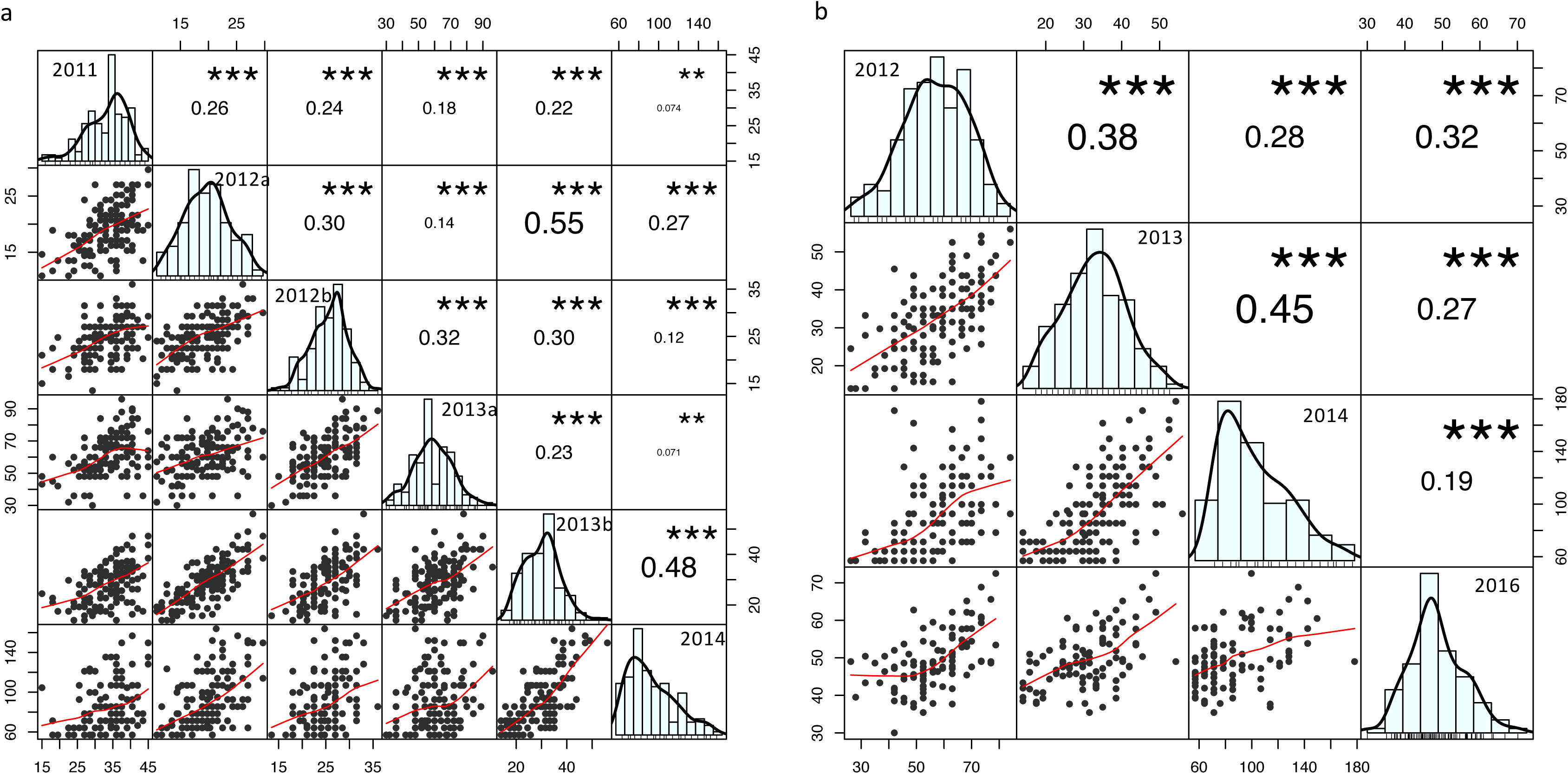
Pearson correlation matrix of powdery mildew area under the disease progression curve phenotype data for the strawberry mapping populations a) ‘Emily’ x ‘Fenella’ and b) ‘Redgauntlet’ x ‘Hapil’. Numbers are R^2^ values.

### Stable and transient QTL are detected in the individual analyses

Five stable QTL (*FaRPa2A*, *FaRPa4A*, *FaRPa5B*, *FaRPa6D2*, *FaRPa7D*) were associated with powdery mildew resistance in more than one phenotyping event and the combined analysis. The focal SNP associated with powdery mildew resistance, representing the stable QTL *FaRPa6D2 was* consistently identified on linkage group 6D in Redgauntlet across three phenotyping events and in the combined analysis. This allele is situated within 9.2 kbp of a putative resistance gene containing an RLK domain on chromosome six of the *F. vesca* genome. Highly significant QTL in ‘Emily’ were identified between 0.3-5.9 Mb on linkage group 1C ‘0’ haplotype in four individual phenotyping events and the combined analysis (Table 2 & Sup. Table 1). QTL on linkage group 1C represent some of the most significant markers associated with mildew resistance found in this study, however the QTL position shifts depending upon the phenotyping event. Six focal SNPs were detected in two or more individual phenotyping events and multiple transient QTL were detected (Supp. Table 1).

**Table 2.**
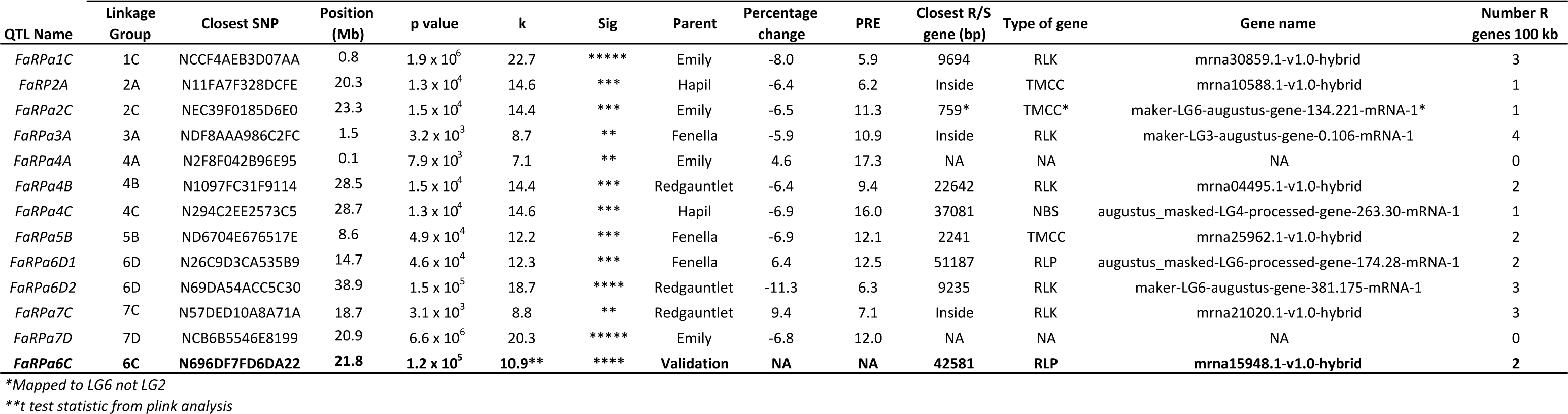
Focal single nucleotide polymorphisms linked with each quantitative trait loci associated with strawberry powdery mildew disease resistance identified through the Kruskal-Wallis analysis using the best linear unbiased prediction calculated across all phenotyping events. Closest resistance gene reported within 100 kbp if applicable. Grey shading indicates alleles that have maintained linkage disequilibrium across the wider germplasm. Bold entries denotes a focal single nucleotide polymorphisms linked with quantitative trait loci associated with strawberry powdery mildew disease resistance identified through the targeted marker association study

In the combined analysis 7 QTL identified in the ‘Emily’ x ‘Fenella’ mapping population (Fig 3) with a combined effect of 45.2 % (Proportional reduction of error (PRE) 82%) whereas the combined effect of the 5 QTL identified in the ‘Redgauntlet’ x ‘Hapil’ mapping population was 40.4% (PRE 45.1%; Fig 4). Of the 12 QTL identified, ten were associated with putative resistance genes in *F. vesca*, three of which fall inside a resistance gene (Table 2). Although none of the identified QTL fall within the assigned threshold of the putative MLO genes, *FaRP2A* is 133,210 bp away from the MLO-like-protein gene *04.XM_004290718.2*. The combined analysis pulled out nine QTL identified in at least one of the individual phenotyping events and three novel QTL. No epistatic interactions were found between QTL identified in the combined analysis, indicating resistance is controlled by additive genetic components in the two populations.

**Fig. 3.**
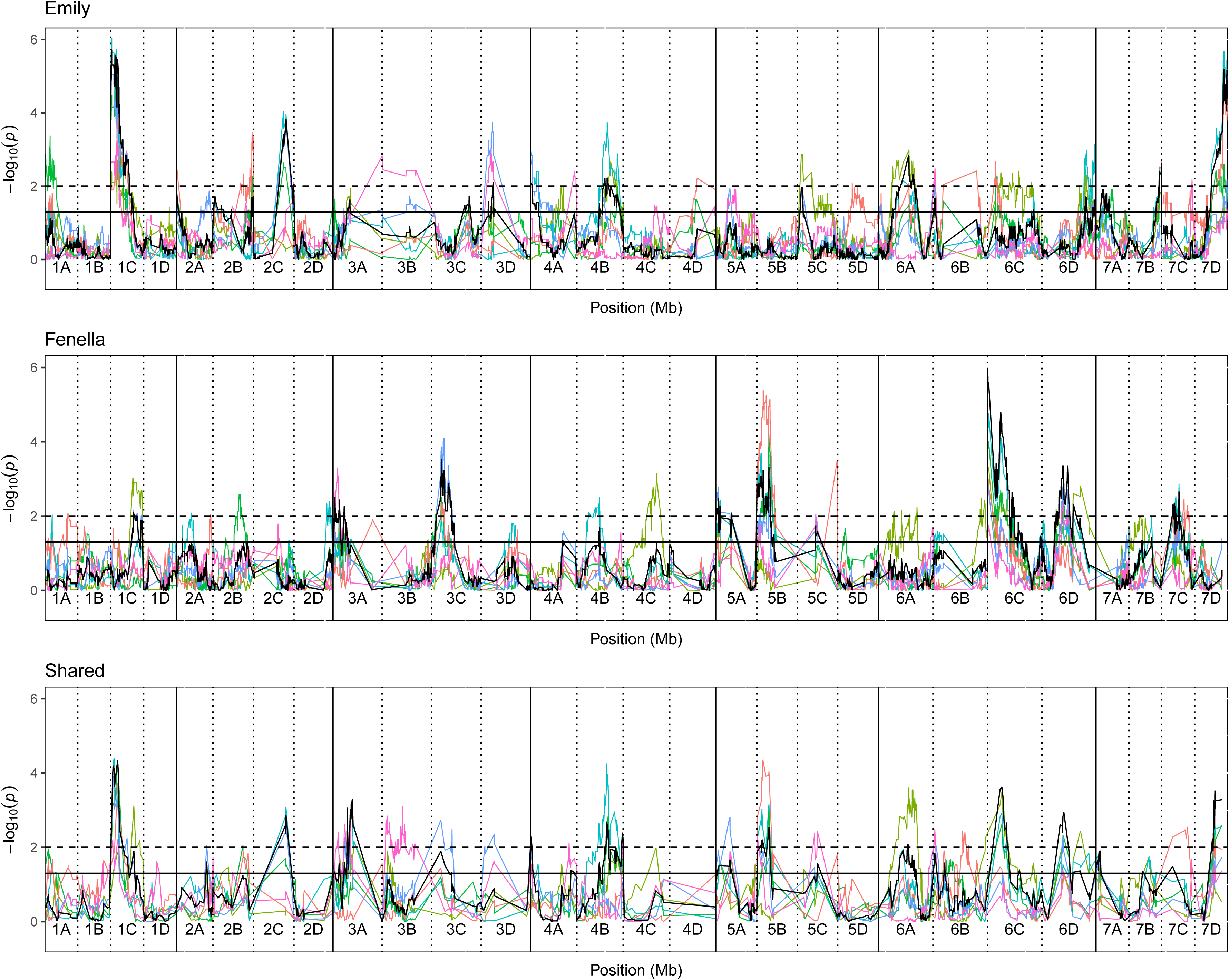
Kruskal-Wallis-log_10_ p-values denoting the association of single nucleotide polymorphism with strawberry powdery mildew disease scores at each position in the octoploid strawberry genome in cM. Panels represent markers segregating in ‘Redgauntlet’, ‘Hapil’ and both parents. Labels 1A-7D denote the 28 linkage groups. Solid horizontal line is *p*= *0.05*, dashed horizontal line is *p*= *0.01*. Black line denotes combined analysis using the best linear unbiased prediction calculated across all phenotyping events. Colour denotes phenotyping event blue-2012, teal-2013, green-2014, pink-2016.

**Fig. 4.**
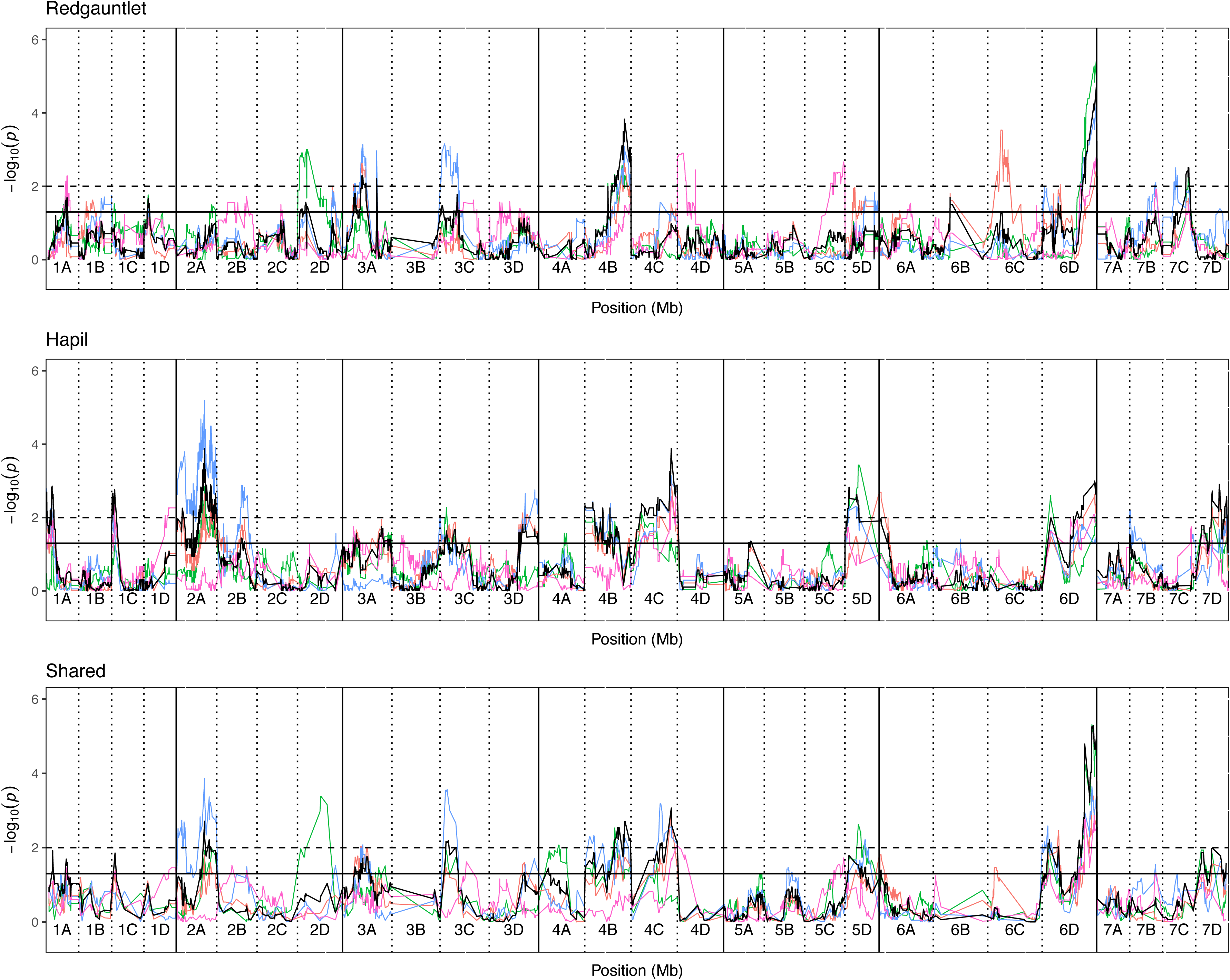
Kruskal-Wallis-log_10_ p-values denoting the association of single nucleotide polymorphism with strawberry powdery mildew disease scores at each position in the octoploid strawberry genome in cM. Panels represent markers segregating in ‘Emily’, ‘Fenella’ and both parents. Labels 1A-7D denote the 28 linkage groups. Solid horizontal line is *p*= *0.05*, dashed horizontal line is *p*= *0.01*. Black line denotes combined analysis using the best linear unbiased prediction calculated across all phenotyping events. Colour denotes phenotyping event olive green-2011, light blue-2012a, green-2012b red-2013a, blue-2013b (Spain), pink-2014.

### Detected QTL explain a large portion of the observed phenotypic variation

The coefficients of determination (R^2^) for the linear regression models of all phenotyping events show positive relationships between predicted and observed values, with between 20.6 % and 73.1 % of variation in observed scores explained by the identified QTL (Table 1).

### Some QTL are detected in similar regions across the two populations

No ‘neighbouring’ focal SNPs within 1.5 Mb were identified in combined analysis’ between the ‘Redgauntlet’ x ‘Hapil’ and ‘Emily’ x ‘Fenella’ populations (Fig 5). The individual analyses identified neighbouring focal SNPs from both populations on linkage groups 3C and 7D (Sup. Fig 1). The neighbouring markers on the top of linkage group 3C are 3.3 cM apart respectively in the ‘Redgauntlet’ x ‘Hapil’ population with the resistance-conferring allele present in the same phase. The neighbouring focal markers on 7D are present on different parents in the ‘Redgauntlet’ x ‘Hapil’ population however no recombination is observed between the markers, therefore we conclude that the QTL identified in the two populations on linkage group 7D represent the same QTL.

**Fig. 5.**
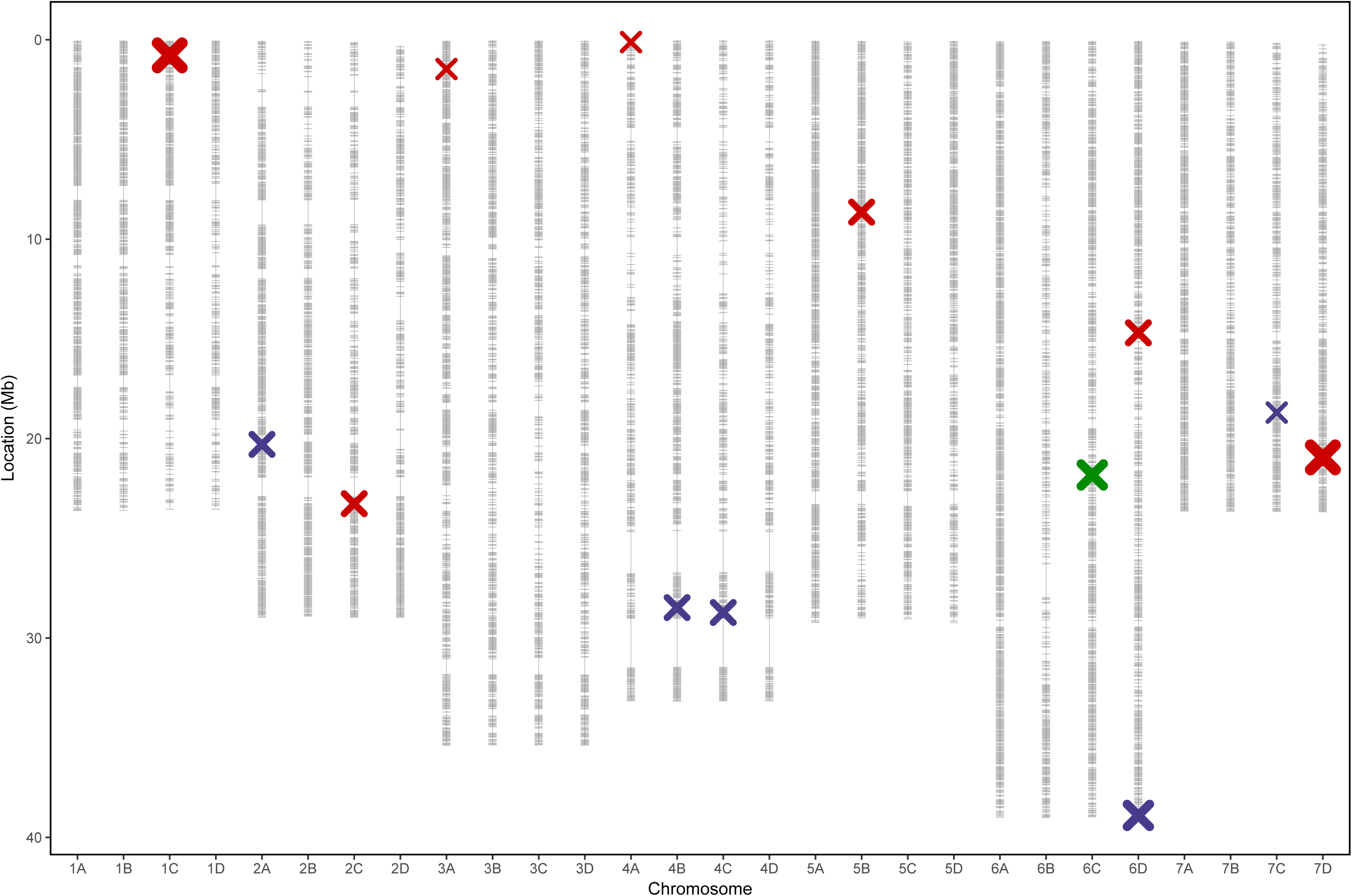
Linkage map displaying 35154 marker positions (grey) in Mb for 28 linkage groups of octoploid strawberry (1A-7D) marker positions scaled to the *F. vesca* genome. QTL locations from combined analysis ‘Emily’ x ‘Fenella’ (red) ‘Redgauntlet’ x ‘Hapil’ (purple) and from the targeted marker association analysis (black) point size denotes significance level of QTL.

### QTL are poorly associated with phenotype in the wider germplasm

The combined transferable QTL analysis for the ‘Emily’ x ‘Fenella’ population produced four i35k substitute SNPs co-localising with focal SNPs identified in the i90k analysis whereas the position of two focal SNPs had shifted and one was not identified (Sup. Fig 2). The combined transferable QTL analysis for the ‘Redgauntlet’ x ‘Hapil’ population produced four i35k focal SNPs co-localising with those identified in the i90k analysis and one focal SNPs locations had shifted. This analysis was associated with a slight loss in power to detect QTL positions but allowed QTL to be screened across the wider germplasm. One of the focal SNPs identified in the combined analysis on linkage group 4C maintained a strong association with resistance across the wider germplasm. This QTL explained 31.6 % of the variation in disease scores observed in the validation germplasm (Sup. Fig 3) The association analysis of the validation set identified multiple SNPs representing a single locus on linkage group 6C (Fig 6; Table 2), however this locus does not appear to contribute to the resistance of the mapping populations.

**Fig. 6.**
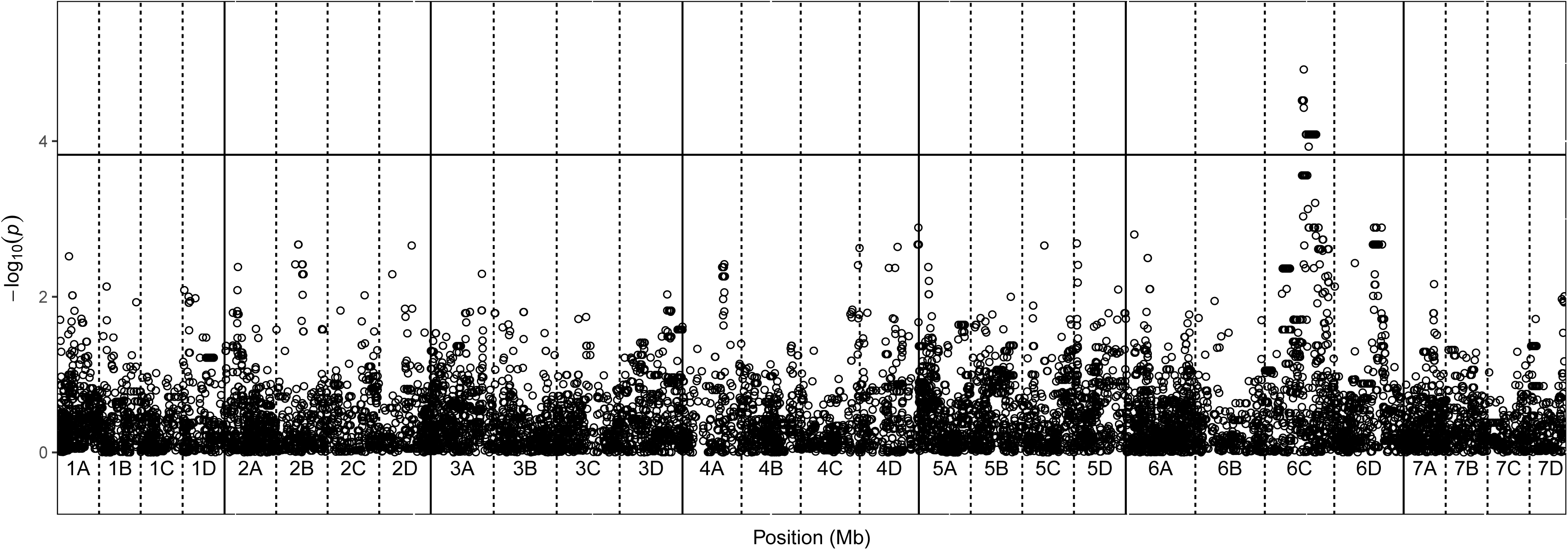
Unadjusted-log_10_ p-values from plink denoting the association of single nucleotide polymorphism with strawberry powdery mildew disease scores at each position in the octoploid strawberry genome in cM, where known. Labels 1A-7D denote the 28 linkage groups. Solid horizontal line is *p*= *0.05* after Benjamin-Hochberg correction.

## Discussion

Comparison of disease scores across phenotyping events revealed the presence of a Genotype x Environment interaction in both populations. This was also observed in similar experiments from other groups (Kennedy et al 2013). The variation in phenotypic scores between each experiment can be explained by differences in a combination of (i) the genetic diversity of the inoculum source (ii) inoculum load and (iii) environmental conditions.

Each phenotyping experiment relied on natural inoculum from nearby plantings resulting in a different admixture of inoculum. There has been no report of race structure between *P. aphanis* and strawberry to date, furthermore isolates from Italy and Israel were found to be homogenous after attempts to develop five discriminatory markers revealed monomorphic loci (Fiamingo et al 2007). It was hypothesised that this may be attributed to low genetic variability in strawberry germplasm or low variation between mildew isolates (Xu et al 2008b). However, there is some evidence to suggest heterogeneity between populations of *P. aphanis:* namely the evolution of fungicide resistance (Sombardier et al 2010) and the resistance breaking in the cultivar Korona, likely due to the evolution of more virulent strains (Davik and Honne 2005). Reports of the production of ascospores in UK infections indicates the presence of a sexual cycle within powdery mildew (Xu et al 2008a), high recombination associated with this life cycle typically leads to greater genetic diversity than observed in asexual reproduction (Barrett et al 2008). Qualitative resistance is associated with race specific interactions and typically the resistance is non-durable due to the R gene targeting a dispensable effector gene (Geiger and Heun 1989; Vleeshouwers et al 2011). It was observed that resistance to *P. aphanis* within new strawberry selections is not durable over time and across varying environmental conditions, however, it cannot be determined whether this is due to unstable resistance or variable mildew strains (McNicol and Gooding 1979; Nelson et al 1995; Xu et al 2008b). These examples indicate the requirement for constant breeding and selection for powdery mildew resistance in strawberry.

It has been suggested that different genes confer resistance to mildew depending on the inoculum level (Nelson et al 1995; Nelson et al 1996; Kennedy et al 2013). Due to the natural inoculation method, phenotyping events varied in inoculum load and such variation may create differential induction of systemic resistance. Future mildew infection experiments could aim to quantify field inoculum levels to qualify resistance genes as effective under low or high inoculum levels. The infection levels of neighbouring plants will influence the inoculum load experienced by a given plant and therefore the resistance status (Hughes et al 1997). However, the randomisation, replication of field experiments and combined analysis should mitigate any issues related to spatial autocorrelation.

Environmental conditions affect the sporulation, germination and establishment of *P. aphanis* conidia. Optimum conditions for germination occur between 75-98 % RH and between 15-25 °C with disease symptoms observed 4 days after germination (Amsalem et al 2006). Expression of quantitative disease resistance is influenced by soil, weather and age of plant material (Geiger and Heun 1989). The plants in each phenotyping event will be exposed to different environmental conditions. However, the two phenotyping events 2012a and 2012b were conducted within 500 meters and therefore have experienced a similar macro-environment and inoculum admixture. The coefficient of determination of 30% between 2012a and 2012b phenotyping events indicates a moderate correlation of disease score.

A strong correlation was observed between strawberry cultivar powdery mildew resistance assessed in the field and polytunnel environments (Gooding et al 1981; Nelson et al 1996; Kennedy et al 2013). Therefore our powdery mildew field assessments should reflect resistance levels exhibited in a polytunnel environment and supports transferability of resistance alleles from field to polytunnel environments. Furthermore plant nutrient status has been found to impact mildew resistant status. Indeed, low calcium levels were associated with weakened mildew resistance response in the cultivar ‘Aroma’ (Palmer 2007). However, fertigation was applied to all plots mitigating the potential for plant nutrient status to impact mildew disease scores.

The QTL analysis was performed individually for each phenotyping event, allowing the identification of both stable and transient QTL. A different suite of QTL from ‘Emily’ x ‘Fenella’ were identified as significant for each of the six phenotyping events indicating that the resistance is indeed complex and quantitative. However, six QTL have support across two or three years of phenotyping. Therefore, relatively few stable QTL are observed alongside multiple transient QTL. Stable and transient QTL were also found to control apple powdery mildew resistance (Calenge and Durel 2006). Stable QTL represent alleles involved in disease resistance on the majority of infection events however transient QTL may represent genes that have an environment specific interaction and as such are important in only some infection events. The large number of transient QTL may be attributed to the variation in inoculum source, inoculum load and environmental conditions between treatments. We found multiple QTL of small to moderate effect control disease resistance in both populations. Likewise, previous studies have found multiple small effect QTL which contribute to powdery mildew resistance in strawberry indicating quantitative resistance with both additive and nonadditive genetic components (Simpson 1987; Nelson et al 1995; Davik and Honne 2005).

Previous studies have shown ‘Hapil’ has an estimated breeding value for mildew resistance of 0.036 (± 0.095) showing almost no genetic component for mildew susceptibility status in this cultivar (Davik and Honne 2005). Here we show several QTL associated with resistance can be identified in the Hapil cultivar indicating that powdery mildew disease status contains a genetic component in this cultivar. Heritability factors observed in the study are lower than the field based disease severity in studies conducted by Nelson et al. (Nelson et al 1995) of 0.7, which indicates a strong genetic component. It has also been found that higher heritability values are associated with high infection levels due to a greater uniform inoculum distribution (Nelson et al 1995). Due to the natural inoculation process there is the potential for a patchy inoculum dispersal across the field site, this may explain relatively low heritability scores.

Genotyping and field phenotyping of the validation accessions revealed that one of the resistance QTL was associated with resistance within the wider strawberry germplasm, indicating a strong candidate marker for further study. However, the moderately low transferability of markers highlights the likelihood for multiple sources of powdery mildew resistance across strawberry germplasm and the need for an enhanced panel of genetic markers that fully represent the diversity present in strawberry germplasm.

When validating QTL identified in a bi-parental cross over the wider germplasm, a reduction in the number of informative loci is anticipated due to the lack of LD between the markers ‘tagging’ resistance in a bi-parental population and the QTL. Based upon the design of the i90k and subsequent arrays, it is likely that these only capture a tiny fraction of the genetic variation, based on the limited discovery panel that was used during the array design (Bassil et al 2015). The focus on the selection of common SNPs for the i35k array also means that low frequency markers present on the i90k are also absent (Verma et al 2017). Furthermore, if QTL are at low frequencies in the wider population, it is unlikely that the underpowered preliminary association study that we have carried out will have sufficient power to detect QTL.

Nonetheless, here we observe one conserved resistance QTL across the wider strawberry germplasm and highlight the potential for some limited transferability of markers into resistance breeding. Future work will seek to identify the candidate resistance genes associated with the QTL on linkage group 1C and 6D and screen for presence of more candidate resistance genes across the wider germplasm. Work will seek to identify the mechanism of resistance in the cultivars ‘Emily’ and ‘Redgauntlet’. Such work has been conducted on the powdery mildew resistant cultivar ‘Aroma’, where poor colony establishment was associated with the identification of a putative antimicrobial protein (Palmer 2007). Additionally, lower conidial attachment was associated with high cutin acid in ‘Aroma’ leaf cuticles, indeed high cutin acid has been extracted from many resistant strawberry cultivars (Peries 1962; Jhooty and McKeen 1965). Relatively high powdery mildew resistance was observed in Florida cultivars from wild accessions of *Fragaria virginiana* after successful introgression into strawberry breeding germplasm highlighting the potential of natural reserves of resistance to be used in crop breeding programmes (Kennedy et al 2013).

The magnitude of disease symptom variation shows great potential to enhance mildew resistance. The most resistant accession was 3.2 times more resistant than the most susceptible within the validation accessions. We conclude that multiple QTL of small effect control disease resistance and that principally a different suite of alleles control resistance in the two studied populations, with a limited overlap. The small effect size of loci promotes a genomic selection breeding approach for powdery mildew resistance in strawberry, as this may be most effective at capturing the wide range of small effect QTL that are likely to be present in a breeding programme. Any training population would need to be phenotyped in multiple environments and years in order to capture the diverse expression of QTL and fully maximise the power of a GS approach. Furthermore, a more detailed study of the pathogen’s population structure and host interactions is needed to quantify the contribution of pathogen diversity to transient QTL.

Ultimately, the production of mildew resistant strawberry cultivars will reduce grower reliance on chemical fungicide for control of powdery mildew; such control options are particularly important with respect to reducing consumer concerns over pesticide residues and also where deregulation of existing fungicide actives is reducing disease management options.

## Acknowledgements

The authors acknowledge funding from the Biotechnology and Biological Sciences Research Council (BBSRC) BB/K017071/1 and Innovate UK project 100875. The authors acknowledge Phil Brain, Eric van de Weg and Xiangming Xu for statistical advice. The authors also acknowledge INRA and CRAG for contributing their mapping population data to the strawberry consensus linkage map.

## Supplementary Figure Legends

**Supplementary Fig 1** Linkage map displaying 35154 marker positions (grey) in Mb for 28 linkage groups of octoploid strawberry (1A-7D) marker positions scaled to *F. vesca* genome. QTL locations from each phenotyping event represented.

**Supplementary Fig 2** Linkage map displaying marker positions (grey) in Mb for 28 linkage groups of octoploid strawberry (1A-7D) marker positions scaled to *F. vesca* genome. Markers overlapping between the validation set and ‘Emily’ x ‘Fenella’ and ‘Redgauntlet’ x ‘Hapil’ populations are red and blue “-” respectively. QTL locations from combined analysis ‘Emily’ x ‘Fenella’ (red) and ‘Redgauntlet’ x ‘Hapil’ (purple).

## Author Contributions

RJH, DWS, DJS - Conceived, designed and analysed experiments.

AJP - Propagated plant material

KJM, NH, KP, ML, AK, HMC, JH - Recorded pathogenicity data in field experiments

RJV and AK – Analysed SNP data and made linkage map.

HMC – QTL mapping and statistical analysis

FW, ML and LA extracted gDNA for SNP chip analysis.

MKS-created R gene database and pipeline used for plink analysis

HMC and RJH wrote the manuscript. It was not possible to contact ML, KJM and LA for approval.

